# Tumor purity adjusted beta values improve biological interpretability of high-dimensional DNA methylation data

**DOI:** 10.1101/2022.03.04.483052

**Authors:** Staaf Johan, Aine Mattias

## Abstract

A common issue affecting DNA methylation analysis in tumor tissue is the presence of a substantial amount of non-tumor methylation signal derived from the surrounding microenvironment. Although approaches for quantifying and correcting for the infiltration component have been proposed previously, we believe these have not fully addressed the issue in a comprehensive and universally applicable way. We present a multi-population framework for adjusting DNA methylation beta values on the Illumina 450/850K platform using generic purity estimates to account for non-tumor signal. Our approach also provides an indirect estimate of the aggregate methylation state of the surrounding normal tissue. Using whole exome sequencing derived purity estimates and Illumina 450K methylation array data generated by The Cancer Genome Atlas project (TCGA), we provide a demonstration of this framework in breast cancer illustrating the effect of beta correction on the aggregate methylation beta value distribution, clustering accuracy, and global methylation profiles.

## Introduction

Epigenetic alterations in tumor cells are a hallmark of cancer, and changes in the DNA methylome represent the first, and to date the best characterized, example of a bona-fide epigenetic mechanism for altering gene regulation in cancer. Understanding epigenetic alterations and their effect on tumorigenesis has therefore long been a research focus in cancer and numerous attempts at capturing epigenotypes of cancer have been made over the years [1–6].

Carcinogenesis is however a complex process involving both epigenetic and genetic insults to the human genome as well as an intricate interplay between the tumor compartment and the surrounding normal tissue as well as cells of the immune system. Consequently, resected bulk tumor samples are not completely pure with respect to tumor cells, but instead consist of a complex mixture of cell types representative of the tumor and its microenvironment. This mixture of malignant and different non-malignant cell types has profound implications on the readout of, e.g., high dimensional genomic profiling techniques. For instance, for somatic mutations occurring in tumor cells the observed variant allele frequency, i.e. the number of sequence reads with a variant divided by the total number of reads covering a specific locus, becomes a function of the proportions of malignant and non-malignant cells. As different cell types also have different epigenetic states it follows that this admixture of cell types also creates a mixture of epigenetic states in the analyzed tissue, making the discovery and characterization of pure epigenetic subtypes of a given cancer type a challenging task.

To circumvent the issue of mixed cell types in bulk tumor specimens and peripheral blood different approaches have been proposed to deconvolve RNA-sequencing [7–9] or global DNA methylation data [10–14] to provide estimates of the magnitude and/or nature of the tumor and non-tumor compartments. These methods have often focused on quantifying the immune component of the tumor compartment and relied on flow-sorted gene expression or DNA methylation data generated from purified blood cell types as the basis for deconvolution.

Another category of methods has instead focused on quantifying the purity of tumor samples from DNA/RNA-sequencing or DNA methylation data without trying to delineate cell types within the non-tumor compartment. Methods for deriving purity estimates from DNA methylation data have been reported by several groups, e.g. [15–18]. With respect to the performance of these methods for estimating sample purity, sequencing-based estimates have generally constituted the golden standard measure of comparison in the respective studies.

Although most developed and used tools for DNA methylation deconvolution and/or tumor purity analysis have been aimed at quantifying tumor purity or the admixture of infiltrating cell types, few have been aimed at correcting for it globally, but have rather proposed using purity as a covariate in group comparisons and/or clustering [17,19–22]. In the case of tools such as those available in the InfinuimPurify-package, which includes a number of approaches directed at controlling for purity in epigenomic analyses, these methods frequently make use of matched reference normal samples for establishing the baseline methylation state of the non-tumor compartment and are currently only implemented on the Illumina Infinuim 450K platform [19]. The requirement of available “normal” baseline data is not always simple to meet and although the Infinium 450K remains the historically most used methylation profiling platform, it has subsequently been replaced by the EPIC 850K array [23].

Accounting for confounding signal derived from the tumor microenvironment (TME) is conceptually straightforward and a theoretical perfect separation is obtainable at CpGs in which the tumor and non-tumor compartment respectively show homogenous and diametrically opposed DNA methylation states. Indeed, this assumption forms the basis for most proposed correction and estimation algorithms and logic dictates that it is only at sites where methylation differs between the tumor and non-tumor compartments that deconvolution strategies can separate alleles contributed by the respective components of the aggregate sample. For the purpose of estimating overall tumor purity this assumption typically only needs to be met for a small minority of the hundreds of thousands of assayed CpG-sites in order for robust estimates to be made. But in order to correct for the TME influence on tumor methylation estimates, one needs to account for the fact that only a minority of tumor cells alter their methylation states in relation to the normal tissue background. This invalidates a simple assumption of one tumor- and one normal compartment with diametrically opposed methylation states, which, e.g., the InfiniumPurify-function implicitly makes [19].

In this study we apply a strategy based on the usage of an estimated tumor cell content, flexible mixture modelling, and linear regression, to correct high-dimensional DNA methylation data at an individual CpG level. The direct basis for this work stems from the observation that CpGs linked to silencing of the well-known *BRCA1* tumor suppressor gene display a two-population CpG methylation pattern in triple negative breast cancer (TNBC, tumors that are negative for estrogen and progesterone receptor expression and lack amplification of the human epidermal receptor growth factor 2/erythroblastic oncogene B, *HER2/ERBB2*, gene) highly correlated with tumor cell content estimated from whole genome sequencing [24]. While *BRCA1* promoter hypermethylation is frequent in TNBC [24] it is well established that somatic hypermethylation underlying, e.g., the *RB1* gene hypermethylation in retinoblastoma [25] or the CIMP-phenotypes of colorectal cancer [1] or glioblastoma multiforme [2] only affects a subset of all tumors of a given cancer type. Thus, when a normal-tumor DNA methylation difference exists, only a minority of tumors display a methylation state that differs from that of the non-tumor compartment. Here, we believe that accounting and correcting for this fact using modelling on an individual CpG level and allowing for the presence of multiple tumor methylation states, results in overall more binary methylation profiles. Such binary profiles may enhance clustering performance of methylation data and characterization of novel tumor suppressor gene loci. As a result of the investigations in global DNA methylation data from breast cancer in this study we also propose that the reliance on normal samples for deconvolution may be overstated as reliable estimates of normal microenvironment methylation states can be obtained from bulk methylation data. We present a proof-of-concept analysis in the TCGA breast cancer (BRCA) cohort, showing that our approach improves the biologically relevant Basal vs non-Basal contrast, results in more binary and thus cleaner methylation profiles in clustering analyses and increases the overall biologically relevant contrast in global and focused analyses. We expect that our observations and approach can be developed further and/or integrated into pre-existing software solutions to improve the capacity of these for de-noising bulk methylation data and allow for more unbiased epigenomic analyses that may yield novel insights into epigenetic phenotypes of cancer and new leads in the search for tumor suppressor genes frequently inactivated by DNA methylation.

## Materials and Methods

### Data sets

We obtained *BRCA1* gene associated CpG methylation data based on Illumina EPIC arrays from Glodzik et al. for 235 TNBC cases together with corresponding *BRCA1* gene pyrosequencing data and tumor purity estimates from WGS (PMID: 31570822) [24,26]. Using the GDC data portal (https://portal.gdc.cancer.gov), we obtained data manifests covering The Cancer Genome Atlas (TCGA) consortium breast cancer (BRCA) cohort with data available on the levels of RNA-sequencing, whole exome sequencing (WES), 450K methylation, as well as copy-number and LOH status. For raw Illumina 450K idat files, we relied on the GDC legacy portal. Data were downloaded using the Genomic Data Commons (GDC) data transfer tool. In a first step platform overlaps were used to filter samples with incomplete data on all levels. In addition, samples were filtered against a previously published data blacklist and were required to have available purity estimates [27], and pass internal quality checks with respect to 450K array normalization. Additionally, PAM50 [28] molecular subtype calls for each sample were obtained from Thorsson et al. [29]. The final cohort consisted of 630 female breast cancer samples. Additionally, custom CpG annotations were derived for the Illumina 450K platform by mapping CpG coordinates to TCGA RNA-seq gene models, CpG density measures as defined by Saxonov et al [30] and Weber et al. [31], TCGA BRCA ATAC-seq peak overlaps [32], repeatMasker-overlaps [33], ENCODE candidate cis-regulatory elements [34], and ENCODE ChIP-seq peak overlaps for 340 transcription factors in 129 cell lines [35]. The complete TCGA data generating workflow is available on github under the following link (https://github.com/StaafLab/tcgaBrca).

### DNA methylation data preprocessing

Raw idat files were processed using the function preprocessFunnorm [36] as implemented in the R-package minfi [37]. Default parameters were used with patient gender estimated using the built-in function getSex in minfi. Infinium 450K probes annotated as poor-performing by Zhou et al. [38] were filtered out, leaving 421 368 unique probes (freeze 2018-09-09). Additionally, a platform-related effect on CpG methylation beta values between the two utilized probe chemistries was adjusted for using the method of Holm et al. [4]. As a normal reference data set, we used GSE67919 [39] downloaded from the Gene Expression Omnibus (GEO) and corrected for the Infinium I/II effect in analyses related to our correction method. For use with the “InfiniumPurify” method (below) we did not perform Infinium I/II correction on normal samples based on the observation that this step influenced the performance of the method negatively (data not shown).

### A strategy to model and correct CpG methylation with respect to tumor purity

Based on the observation that DNA methylation and tumor purity often interact to produce mixed methylation states that can be modelled by one or more linear functions we devised an algorithm in the R programming environment [40] to correct for this effect that produces purified tumor-as well as inferred “normal” methylation profiles for a cohort of samples to be corrected. The first objective of the algorithm is to on a CpG basis identify the presence of up to three natural populations of samples that arise as a product of the admixture of tumor and non-tumor cells, and then use the discovered populations in a second step to derive estimates of the purified tumor and normal background methylation states. Briefly, the algorithm uses the DNA methylation of a single CpG at a time for a given set of samples as well as a matched global purity estimate variable for each sample as input. A small amount of gaussian noise N(0,0.005) is added to each sample’s methylation estimate to counteract the effect of zero standard deviation (SD) populations on modeling. The function “stepFlexmix” in the FlexMix package for model-based clustering [41] is then applied with methylation as the dependent and 1-purity as the independent variables and with parameters K=1-3 and nrep=3. A best fit model is chosen using the function “getModel” with “BIC” as criterion and the clusters (N=1-3) are obtained using the function “clusters”. Next, a linear model is fitted to each identified population using the original methylation estimate as the dependent and 1-purity as the independent variable to obtain a linear fit with the intercept serving as the pure tumor methylation state. To obtain the normal background methylation state for each population the same regression is performed with purity as the independent variable. The final methylation state is in each case formed by adding the residuals of each fit to the obtained intercept(s). In a final step, values <0 and >1 are set to zero and one, respectively. The individually adjusted CpG methylation estimates for each sample are then aggregated to reform a data matrix of the same dimensions as the input beta matrix (CpGs as rows, samples as columns). For improved run speed the function is implemented using the R-package “parallel” so that calculations for separate CpGs are distributed across a user-specified number of available cores. To produce deterministic results, each individual CpG is also paired with a unique RNG seed number that makes each parallelized operation fully reproducible. Scripts required to perform beta adjustment are available on github under the following link (https://github.com/StaafLab/adjustBetas).

### InfiniumPurify 450K methylation analysis

The InfiniumPurify R package [19] contains functions for both the estimation of tumor purity (function getPurity) and correction of beta values (function InfiniumPurify). The correction function requires a tumor set as well as a normal data set (both N>20) in combination with a purity estimate for each tumor sample. For all runs we used the 630 TCGA BRCA samples processed as described above. Normal methylation profiles were obtained from GSE67919 and were not corrected for Infinium I/II effects when used with InfiniumPurify. The DNA sequencing-based purity variable was obtained from [27].

## Results

### Modeling DNA methylation as a function of tumor purity

We previously analyzed genome-wide DNA methylation patterns in the context of *BRCA1*-mutated vs hypermethylated triple negative breast cancer (TNBC) [24]. As part of this study, we noted anecdotally that CpG loci display different methylation characteristics with respect to tumor purity. An example of one of these patterns is illustrated by the methylation state of a CpG in the *BRCA1*-promoter region in relation to tumor fraction estimates derived from WGS (Figure 1a). The observed methylation pattern across the 235 TNBC samples shows the presence of two distinct populations with diametrically opposed methylation states when accounting for tumor purity. Interestingly, these two observed populations of tumors (hypermethylated and non-methylated) are well described by a combination of linear functions and are well captured using an unsupervised flexible mixture modeling approach that allows for the detection of up to three sample populations (Figure 1b). Although all three populations in Figure 1b would result in equivalent final estimates on the sample level, population 3 (green) can be regarded as redundant and is the result of the mixture modeling algorithm being overly sensitive to minor differences in the data variance structure. This effect can be mitigated by, e.g., adding a small component of normally distributed noise to the methylation estimate, yielding two rather than three populations given the same input variables (Figure 1c). By using the intercept terms of the respective linear regression models for the two populations, as well as purity and 1-purity as the independent variable, an estimate of the methylation state of both the “normal background” (Figure 1d–e) and pure tumor can be obtained (Figure 1f–g) using the equation of the straight line. By then de-trending the respective population beta estimates with respect to 1-purity, a markedly improved separation can be observed between the two tumor populations (Figure 1g). We similarly note that concordant estimates of the methylation state of the aggregate tumor microenvironment (presumably normal breast tissue) are obtained when modeling methylation as a function of tumor purity (Figure 1e).

**Figure 1.**
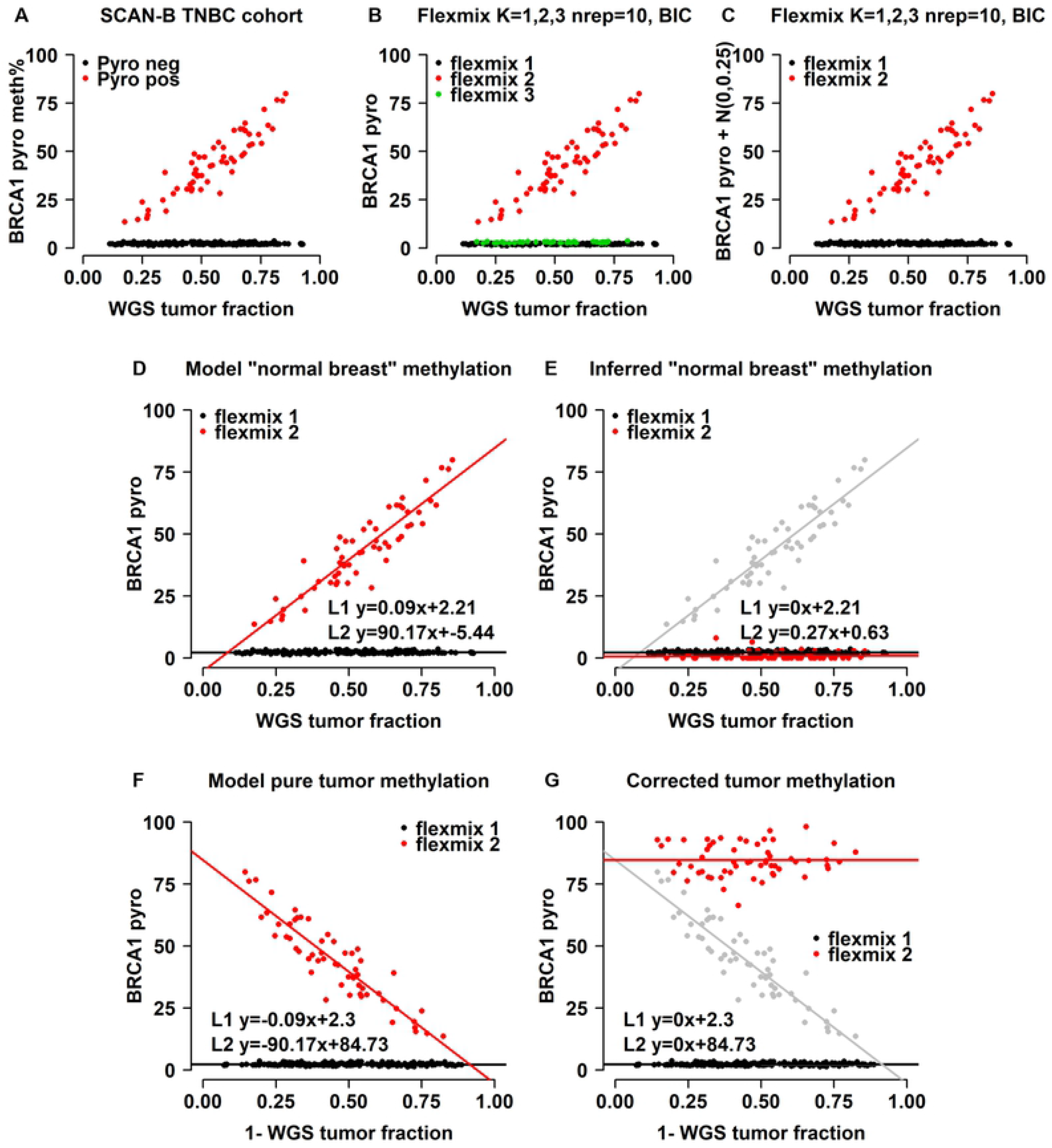
Linear relationship of *BRCA1* promoter methylation percentage with tumor cell content. **A)** Pyrosequencing (pyro) results for *BRCA1* promoter methylation in the SCAN-B 235-sample TNBC cohort plotted against tumor purity derived from WGS. Samples called as *BRCA1* hypermethylated by pyrosequencing are highlighted in red. **B)** FlexMix automated population calls based on linear mixture model with max K=3. FlexMix correctly separates methylated from unmethylated samples but is too sensitive to low-level variance in unmethylated samples. **C)** FlexMix automated population calls based on linear mixture model with max K=3 and a small amount of stochastic noise added to the pyrosequencing data. FlexMix maintains the ability to correctly separate methylated from unmethylated samples after the addition of stochastic noise but less sensitive to spurious noise in the low-variance compartment. **D-E)** FlexMix line fits to data from (C) to extrapolate methylation state of the non-tumor compartment, original population 2 indicated in gray in panel (E). The two line intercepts (L1, L2) identify the correct background methylation state. **F-G)** FlexMix line fits to data from (C) to extrapolate methylation state of the tumor compartment(s), original population 2 indicated in gray in panel (G). The two line intercepts (L1, L2) form the baseline methylation state for the two detected tumor populations.

### A multi-population approach for correcting DNA methylation beta values using tumor purity estimates

Based on the observation that the tumor compartment can display a methylation pattern in which only a subset of tumors diverges from the presumed somatic methylation state, we set out to define a framework for correcting DNA methylation beta values that allows for the presence of multiple separate methylation states in the tumor compartment, as well as estimation of the aggregate methylation state of the non-tumor compartment without the requirement of a-priori information about the normal tissue methylation state or paired normal data. For this we settled on an unsupervised approach using mixture modeling, in which the FlexMix framework is applied on the level of individual CpGs for automated population discovery of samples in a cohort with similar underlying methylation states confounded by non-tumor methylation using a “purity” estimate as a regression covariate.

Our framework is agnostic to how the tumor purity estimate for a sample is obtained. Thus, estimates may be obtained from analysis of genetic data by WES, WGS or SNP microarrays, epigenetic data (e.g., DNA methylation arrays), or pathology estimates. The framework does however intuitively require a subset of cases in a sample cohort to diverge from the presumed somatic methylation state for the population identification to be robust. Based on empirical assessment, and to limit runtimes, we chose a maximum of three populations to model in each iteration (CpG). The FlexMix output is parsed to obtain population designations for each sample and line fits including slopes and intercept terms are obtained for the N discovered populations. If less than two populations are defined, correction for tumor purity is performed by treating all samples as a single population similar to the method implemented in the InfiniumPurify R-package [19]. To improve runtimes, we implemented our method using a parallel computing approach and each CpG is paired with a unique seed number to ensure reproducibility.

### Application of the multi-population method to random CpGs enhances the Basal-Luminal distinction in breast cancer DNA methylation data

To evaluate the performance of purity-corrected methylation data, we extracted 100 data sets of 500 random CpGs (with beta variance >0) each from the Illumina 450K platform for 630 breast cancers from the TCGA consortium and performed agglomerative hierarchical clustering (Pearson distance, Ward’s linkage method) on unadjusted and adjusted data respectively (Figure 2a). We evaluated the performance of beta correction by testing how well the top-level split captured the RNA-sequencing derived PAM50 Basal vs Luminal (non-Basal) tumor assignments. For comparison, we also performed the same clustering using dichotomized beta values (beta >0.3) as described in Hoadley et al. 2018 [27], and beta values adjusted using the “InfiniumPurify” function as implemented in the eponymous R-package [20]. Compared to unadjusted beta values we found increased discrimination of the Basal vs Luminal split in terms of specificity and accuracy using our approach, while beta dichotomization did not improve the discriminatory power neither relative to unadjusted data nor in absolute terms (Figures 2b–c). Using InfiniumPurify yielded better results than dichotomization in terms of most assayed metrics but did not outperform our proposed method.

**Figure 2.**
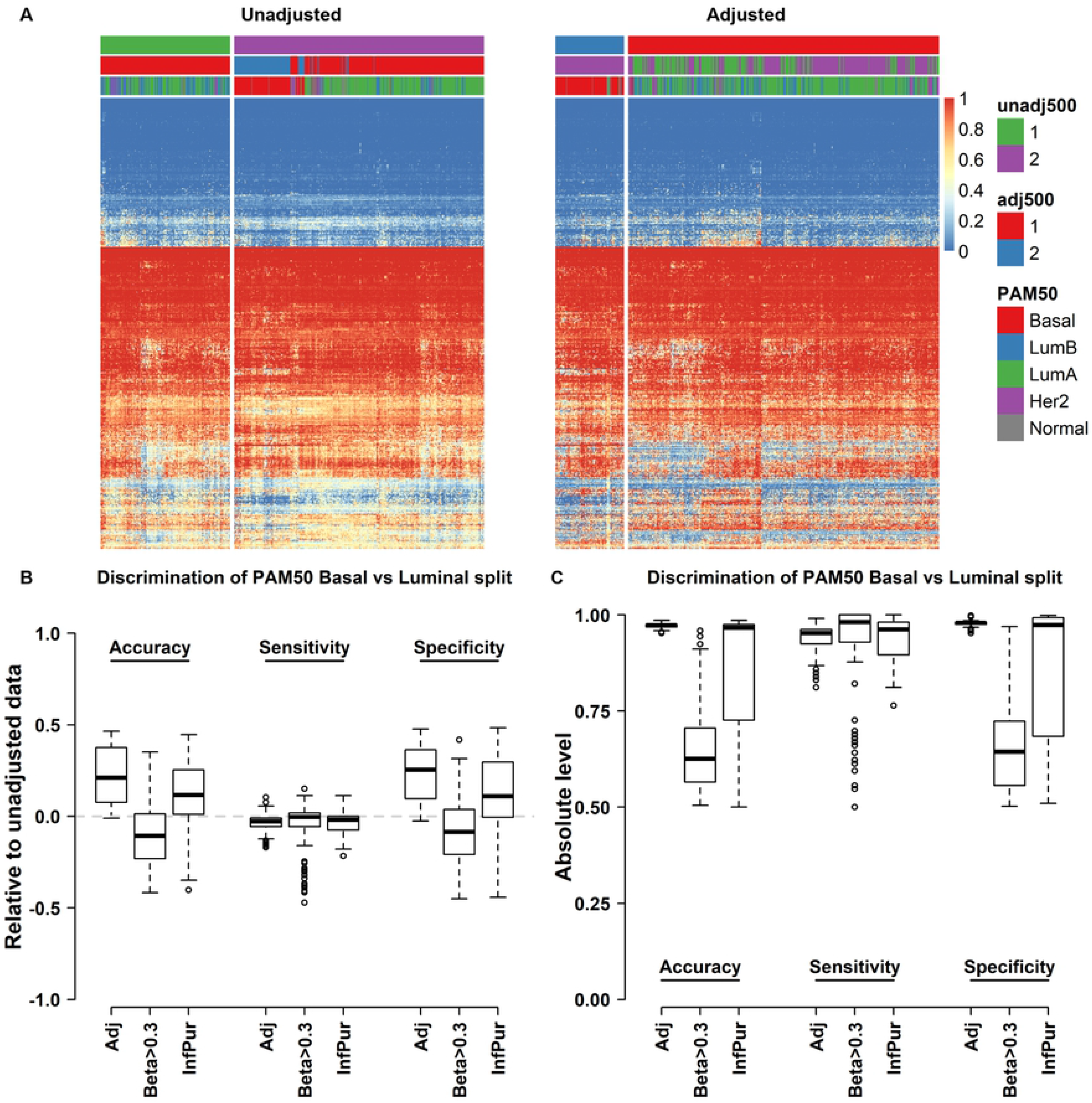
Beta-value correction improves PAM50 Basal vs non-Basal division of TCGA breast cancer samples. Example of hierarchical clustering heatmap of 500 random CpGs from 630 breast cancers from the TCGA consortium before **(A)** and after **(B)** adjustment for tumor purity. Adjusted clustering displays better separation of Basal and non-Basal samples (hierarchical tree cut at K=2). **C-D)** FlexMix adjustment of methylation beta values improves the separation of PAM50 Basal vs non-Basal samples in hierarchical clustering analysis in the TCGA cohort relative **(C)** and absolute **(D)** terms and when compared with dichotomization or InfiniumPurify prior to clustering.

### Deriving corrected beta values and an aggregate estimate of the methylation state of the non-tumor compartment from bulk breast cancer data

To more comprehensively evaluate our approach we applied our method to the top 5000 most varying CpGs across the Illumina 450K platform for the 630 TCGA breast cancer cases. We first evaluated the beta correction using heatmap visualization of uncorrected beta values (Figure 3a middle panel), inferred normal beta values (Figure 3a left), and adjusted beta values (Figure 3a right) using sample and row ordering based on agglomerative clustering of corrected beta values and by selecting a five-group split of samples as the unit of comparison.

**Figure 3.**
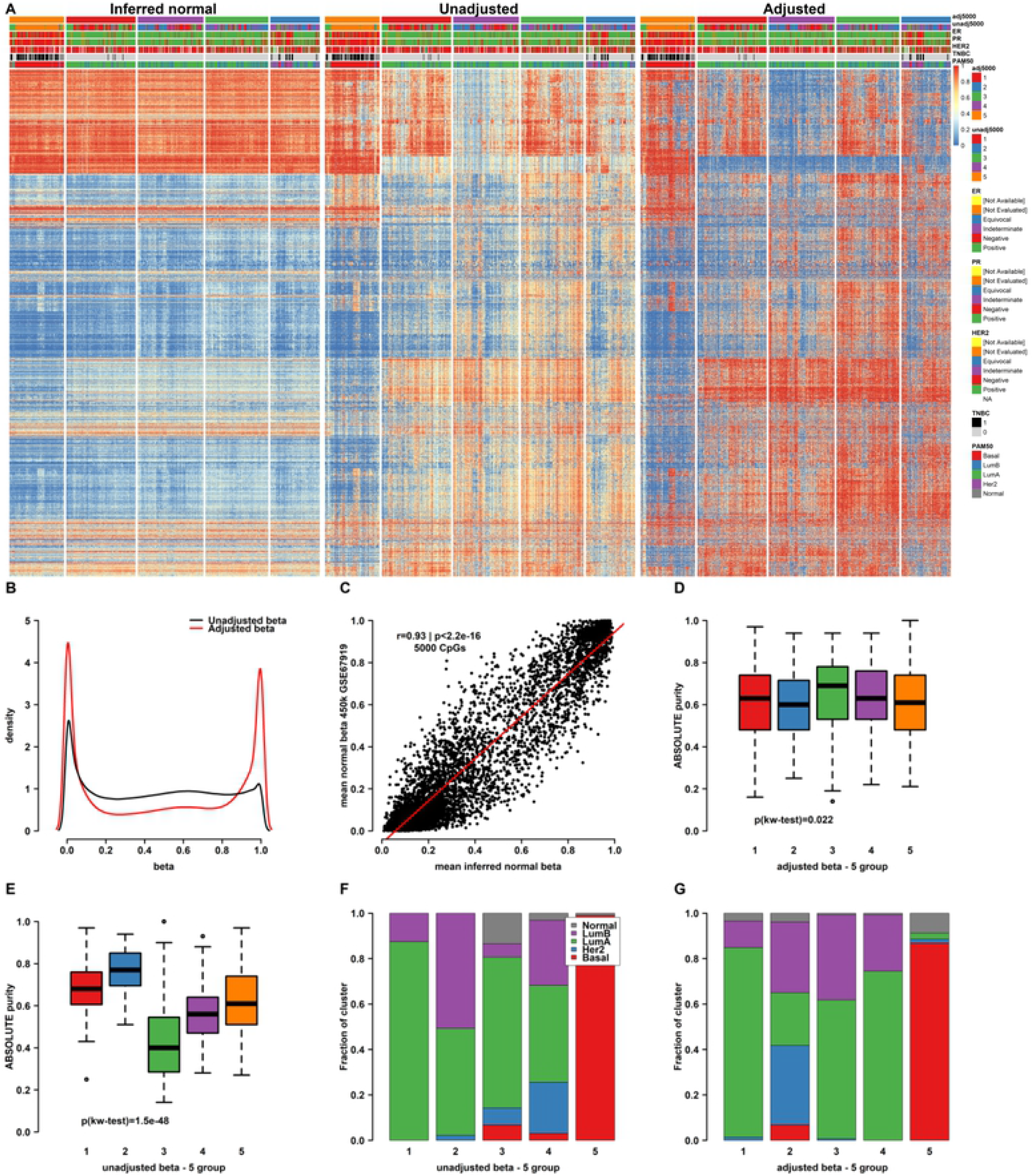
Purity adjustment and the methylation state of the non-tumor compartment from bulk breast cancer data. **A)** Heatmap clustering of the top 5000 most varying CpGs in the TCGA 630-sample BRCA cohort. Original methylation data (middle) with inferred non-tumor methylation (left) and purity-adjusted (right) data presented for comparison and contrast. **B)** Density plot showing effect of purity-adjustment on the beta-value distribution of the top 5000 most varying CpGs. **C)** Scatterplot of mean inferred beta versus mean of “normal breast” methylation beta values for the same 5000 CpG showing excellent overall correlation. **D)** Boxplot visualization of the profound effect of tumor purity when cutting a hierarchical clustering tree on the level of five clusters in unadjusted data. **E)** Boxplot visualization of the much-reduced effect of tumor purity on clustering in purity-adjusted data. **F-G)** Barplot visualization of the distribution of PAM50 subtypes by cluster in unadjusted **(F)** and adjusted **(G)** data showing more discrete partitioning of gene expression subtypes in adjusted clusters.

We also evaluated the aggregate beta distribution in uncorrected and corrected data respectively and found that adjusted beta values were shifted towards the extremes of the beta distribution in comparison with unadjusted beta values consistent with the conception of DNA methylation being binary (Figure 3b). To evaluate the performance of our method for inferring the normal methylation state from analysis of bulk tumor specimens we compared the inferred normal state data from the 630 TCGA breast cancer samples against a public data set of 81 healthy breast tissue samples profiled on the same Illumina methylation platform (GSE67919) [39]. Plotting the mean inferred beta values in our cohort against mean observed beta values in the normal breast samples for all CpGs (5000 most varying) showed a good overall agreement (Pearson’s r=0.93, p<0.001, Figure 3c). We also calculated the correlation coefficients between each individual inferred normal sample and the average methylation state of the same normal breast tissue CpGs and contrasted these with correlations calculated from the corresponding tumor estimates before and after purity adjustment (Supplementary Figure 1). This analysis showed a similar performance on the level of individual inferred normal samples (median r=0.86) as that observed in aggregate. More importantly, it demonstrated a low correlation between normal breast and adjusted tumor methylation states across the top 5000 most varying CpGs in our cohort (median r=−0.04). Moreover, calculating the individual correlation coefficients between unadjusted beta values and mean normal methylation showed a median correlation of 0.4, implying a high overall effect of the TME on native beta values.

Examination of the respective five-group sample sets in terms of tumor purity showed that while tumor purity estimates were a dominant feature differentiating subgroups in uncorrected data (Figure 3d), this effect was largely attenuated in the subgroups defined using corrected beta values (Figure 3e). In terms of co-clustering, while a nearly pure PAM50 Basal cluster was observed in unadjusted data, several Basal tumors co-clustered with luminal and PAM50 HER2-enriched tumors (clusters 4 and 5 in uncorrected data, Figure 3f). In corrected data, a larger fraction of samples classified as PAM50 Basal on the level of RNA-sequencing formed one cluster (cluster 5) in corrected data (Figure 3g). For comparison with the InfinimPuirfy-method, we also performed beta adjustment using the eponymous function as implemented in Qin et al. [19]. Overall, the InfiniumPurify method only seemed to produce modest changes in appearance when visualized as heatmaps (Supplementary Figure 2).

### The effects of purity adjustment on X-chromosome methylation in an all-female cohort

The methylation state of the X-chromosome is unique in that it undergoes lyonization (Xi) in females to achieve dose compensation for an extra gene copy in comparison to the male genome [42]. This yields a trimodal expected methylation pattern at high CpG density positions in females with expected peaks at beta values of 0, 0.5, and 1 respectively. Given that our method produces valid estimates of the normal sample methylome, we expect these estimates to mirror those seen in bona fide normal samples and adhere to the beta distribution expected from biological theory. We therefore extracted CpGs mapping to X-chromosome promoters (N=2921) from both the inferred normal methylomes and GSE67919. For both data sources we plotted the median beta distribution and calculated the fractions falling into the three theoretical X-methylation categories (hypo, Xi, and hyper, Figure 4a). With respect to the three methylation bins the concordance was 90.9%, and with a majority of X promoter CpGs being in a hemimethylated (Xi) state. A minority of CpGs in both data sources were hypomethylated and a subset of these are expected to represent promoters of genes that escape Xi. As a further test of biological coherence, we tested whether the hypomethylated promoters resided in promoters of genes that escape Xi. For this we cross-referenced the gene symbols annotations of the CpGs with a list of 75 genes that were found to escape Xi by Katisr and Linal [43]. As expected, the hypomethylated bin was enriched for localization in promoters of genes escaping Xi (4.6% of hypo CpGs vs 0.8% and 0.6% for Xi and hyper CpGs, respectively). Similarly, plotting the beta distribution of all CpGs mapping to high CpG density positions (N=6736) in adjusted and unadjusted tumor samples showed that purity-adjusted beta values adhere better to the theoretical expectation (Figure 4b).

**Figure 4.**
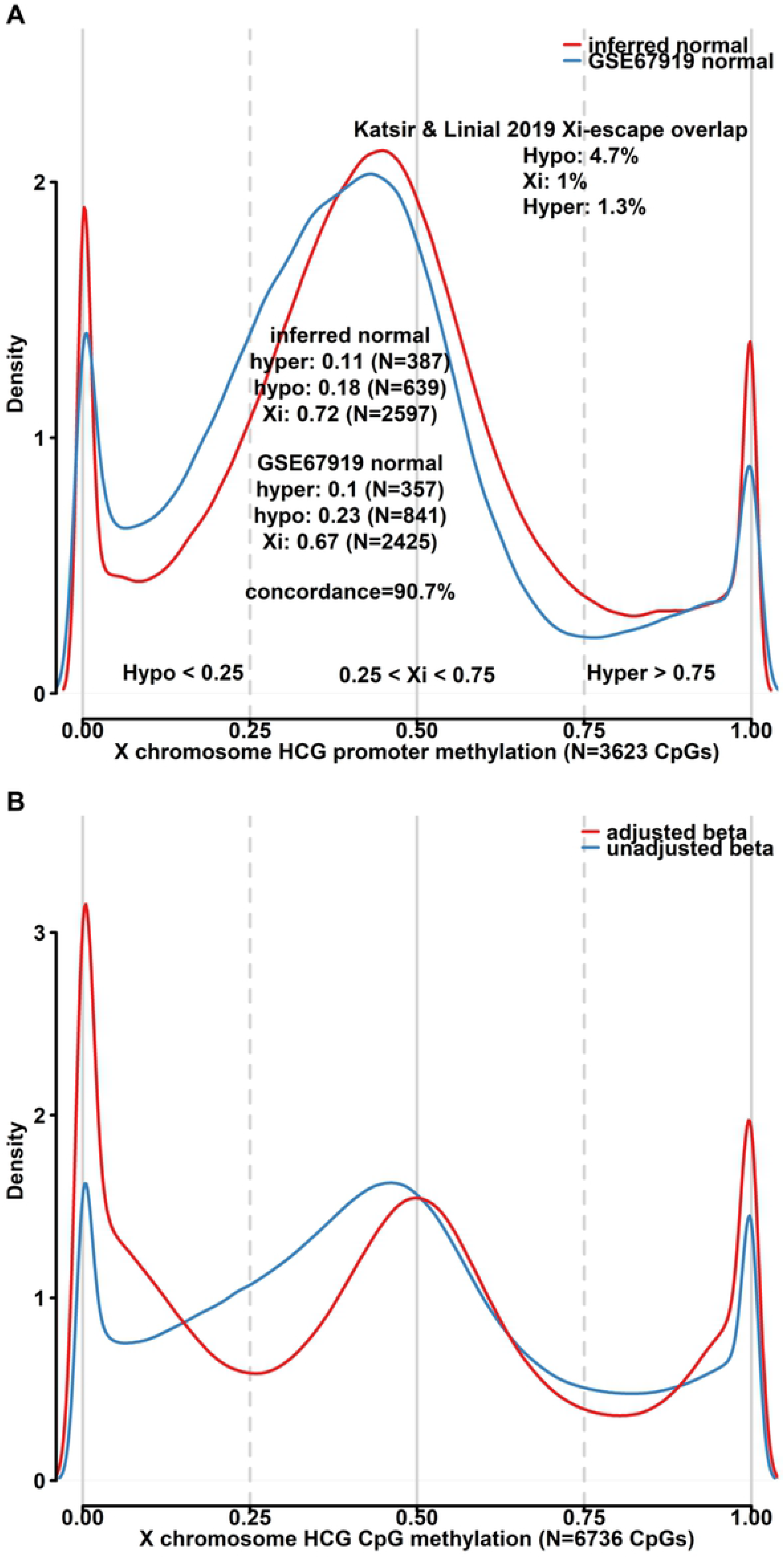
The effects of purity adjustment on X-chromosome methylation. **A)** Density plot highlighting the beta distribution of inferred non-tumor samples (red) and normal breast tissue (blue) in X-chromosome promoters for samples in the TCGA BRCA cohort. Partitioning of promoters based on average methylation state into hypo-, hyper-, and Xi showed high concordance in the respective data sets. Hypomethylated promoters were more frequently found to overlap previously published genes that undergo X-escape. **B)** Purity-adjustment of tumor sample methylation beta values result in a distribution that more closely adheres to the distribution expected from the process of Lyonization.

### The effects of purity adjustment on between-sample contrast at biologically significant classes of CpGs

To establish further biological validity for our beta correction approach, we analyzed pre- and post-adjustment beta values for the top 5000 most varying CpGs from the perspective of native genomic context, and with respect to the 5-group hierarchical clusters, PAM50 subtypes, and TNBC vs non-TNBC cancers.

It has long been known that regions of the genome marked by the Polycomb repressive complex (PRC) in embryonic stem cells are frequently methylated in cancer [44]. Using ENCODE cell-line ChIP-seq data we selected CpGs with the capacity to be marked by the combination of EZH2 and SUZ12 (N=881) among the 5000 top varying CpGs (Figure 5 a). As expected, these CpGs were situated in a CpG-island (CGI) and promoter context and often overlapped TCGA BRCA ATAC-seq peaks. The Basal/TNBC-enriched cluster 5 displayed the lowest aggregate methylation of PRC-bound CpGs while cluster 3 with the highest proportion of Luminal B tumors showed the highest (Figure 5b). The effect of beta adjustment was least prominent in Basal/TNBC- as compared to Luminal tumors as the Basal cancers display the same aggregate methylation state at PRC-marked loci as normal breast tissue (Figures 3a and 5a). Luminal B and HER2-enriched tumors on the other hand more frequently display hypermethylation of PRC-marked loci which is diluted by the presence of non-tumor cells.

**Figure 5.**
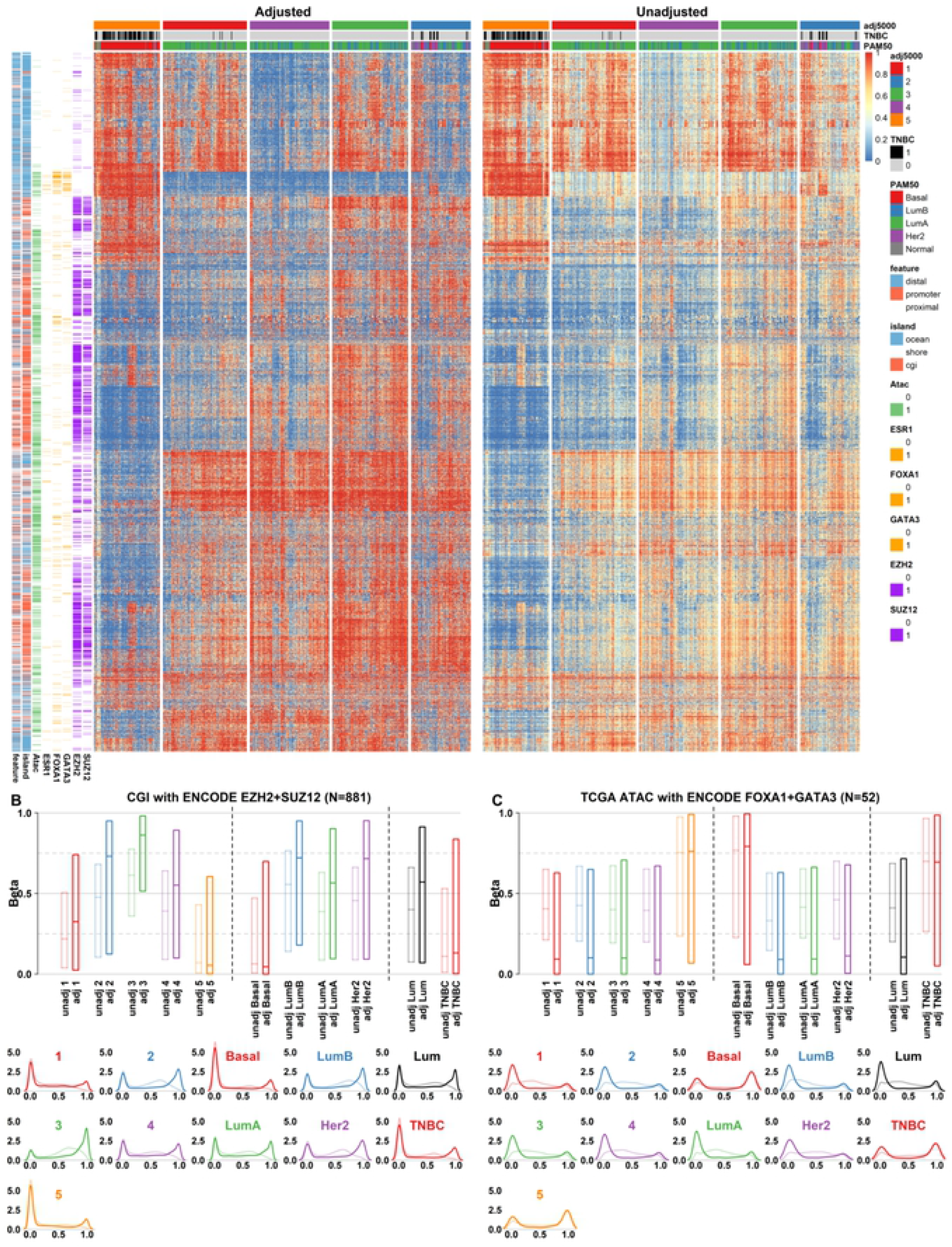
The effects of purity adjustment on between-sample contrast at biologically significant classes of CpGs. **A)** Clustered heatmap visualization of the 5000 most varying CpGs in the TCGA BRCA data set in purity-adjusted (left) and unadjusted (right) data. Row annotation bars highlight genomic features associated with each CpG site and include; genic context, CpG island/shore/ocean localization, TCGA ATAC-seq peak overlap, ENCODE TFBS overlaps for ESR1, FOXA1, GATA3, EZH2, and SUZ12. Samples are clustered based on purity-adjusted data. **B-C)** Boxplot visualization (top) of the beta-value distribution before and after purity-adjustment in the five hierarchical clustering subgroups (left), stratified by PAM50 subtypes (middle), stratified by TNBC-status (right). Below are the same data shown as density plots. Panel **B)** highlights the beta distribution of CGI CpGs with EZH2 and SUZ12 TFBS overlaps. Panel **C)** highlights the same for TCGA BRCA-specific ATAC-seq peaks with overlapping FOXA1 and GATA3 TFBS.

Other prominent transcription factors in BRCA include the pioneer factor *FOXA1* as well as *GATA3* that cooperate with and modulate *ESR1* activity in luminal tumors [45]. To gauge the methylation state of this class of regulatory CpGs, we mapped ENOCDE *FOXA1* and *GATA3* bound sites with overlapping ATAC-seq peaks in TCGA BRCA data (Supplementary Figure 3). A small number of sites (N=52) among the top 5000 CpGs were annotated as having overlaps with all three features. These sites mapped to distal (>5kb from nearest promoter) and ocean designated parts of the genome, consistent with these sites acting as distal enhancers. This class of CpGs were densely methylated in Basal/TNBC tumors as well as inferred normal breast tissue (Figure 5a and 3a respectively) and showed near-universal demethylation in non-Basal tumors. Again, the contrast between pre- and post-correction betas showed considerably improved separation of, e.g., Basal vs Luminal tumors, and highlights that non-tumor tissue again acts to lessen this contrast in unadjusted data.

Finally, we chose to focus on distal CpGs without overlapping ENCODE transcription factor binding sites (TFBS) (N=476) as representative of non-regulatory DNA methylation (Figure 5d). The CpGs were situated in a low CpG density context and did not overlap TCGA ATAC peaks. The methylation state of these CpGs showed a lower association with PAM50 subtypes or a TNBC designation, but instead had more of a gradient-like methylation profile ranging from very low aggregate levels in cluster 4 (luminal A/B) tumors to generally high in cluster 3 (luminal A/B) and 5 (Basal/TNBC) tumors. Generally, demethylation of low-density CpGs has been attributed to lack-of-maintenance or proliferation-related processes, however this cannot likely be the cause in this instance as both HER2-enriched and Basal-like tumors displayed relatively higher methylation levels. In general, Luminal B tumors tended towards lowest overall methylation levels at “non-functional” CpGs, although this was in no way defining for the subtype overall.

### Stability of corrected beta-values to perturbations in tumor purity estimates

In order to quantify the effect of uncertainty in the purity estimate on beta adjustments, we performed a simulation experiment in which purity estimates were perturbed with increasing amounts of normally distributed noise (N(0,s), s=0.01 to 0.19 in increments of 0.02). Beta adjustments were made for the top 5000 most varying CpGs by standard deviation and keeping cohort size (N=630) and all other parameters unchanged. Similarities with unperturbed results were compared for: i) FlexMix population calls on the CpG level, ii) sample-level beta correlation, and iii) correlation of individual inferred normal samples to mean methylation of GSE67919 normal samples. As expected, increasing amounts of noise in purity estimates negatively affect all our calculated metrics proportionally to the magnitude of added noise (Figure 6a). As a rule, the inferred normal estimates had the fastest decay rate with increasing noise, although this effect was not as pronounced on the level of the cohort average as for individual estimates. Similarly, but to a lower extent, FlexMix group calls on the level of individual CpGs were also affected by noise but overall concordance with calls obtained using unperturbed purity estimates remained consistently high (lowest median concordance ~0.9). In terms of sample-level adjusted beta values, a decrease in median correlation and increase in correlation variability was observed but with above 0.95 median correlation in the highest perturbation comparison. Overall, we found that our approach for beta adjustment appeared robust to perturbations of at least moderate magnitudes (mean absolute purity shift in highest noise test ~0.15). Additionally, we tested purity adjustment in the 235 TNBC samples with available WGS purity estimates, Infinium 850K methylation data as well as gold standard pyrosequencing calls for *BRCA1* promoter methylation [24]. The most varying 850K probe in the *BRCA1* promoter showed excellent overall agreement with pyrosequencing-based *BRCA1* hypermethylation calls and applying the FlexMix-based adjustment framework to this single CpG produced highly concordant results (Figure 6b). Analogously to the first perturbation test, we evaluated the influence of increasing the noise level in the purity estimate, this time to the *BRCA1* promoter CpG only. We performed 50 iterations each of beta adjustment across 26 increasingly confounded purity estimates (noise N(0,s), s=0.02 to 0.5 in increments of 0.02). Somewhat surprisingly, overall accuracy remained stable across the tested span albeit with a tendency towards deterioration at higher noise levels (Figure 6c). Interestingly, adding higher levels of pure noise to the purity estimate results in a lower overall magnitude of adjustment, i.e., beta estimates are shifted less on average the more deteriorated the purity estimate is (Figure 6d). With respect to the high maintained accuracy across noise levels, this is a byproduct of only purity estimates being confounded by noise while keeping methylation levels unchanged. The FlexMix framework is therefore still able to reliably separate tumors with differing methylation states, but with increasingly poor performance in finding the correct intercept for the tumor and normal populations as indicated in Figure 6a.

**Figure 6.**
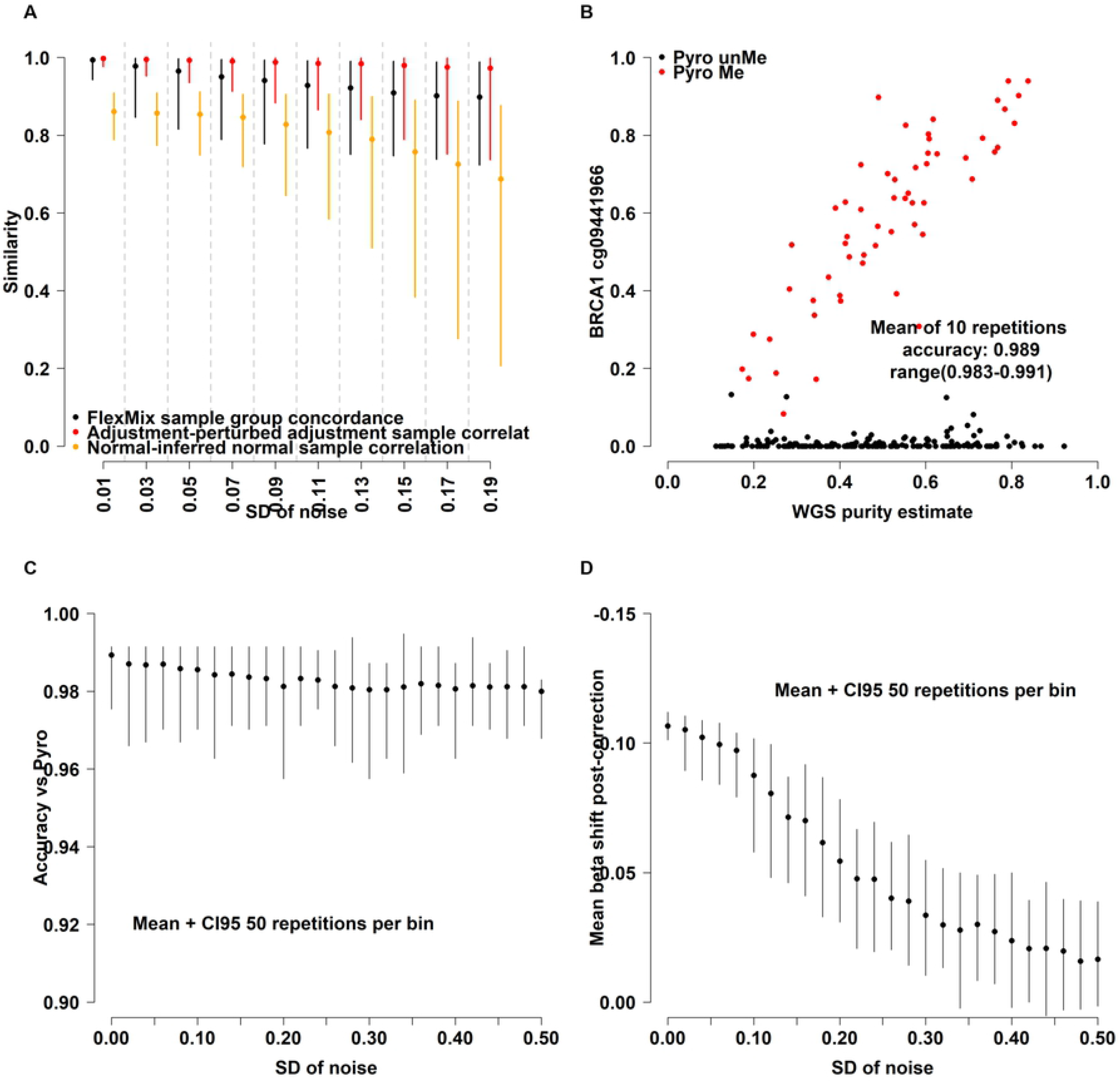
Stability of corrected beta-values to perturbations in tumor purity estimates. **A)** Effect on top 5000 CpGs of increasing amount of noise added to the purity variable prior to beta adjustment as measured by CpG-level concordance of FlexMix sample population calls (N= 5000 CpGs, black), sample-wise Pearson correlation (N=630 pairs, red), correlation between inferred normal breast methylation (iNorm, N=630 samples) and average methylation of true normal breast tissue (Norm, GSE67919). Data presented as median across replications (point) with 95% empirical CI. **B)** Scatterplot showing the most varying 850K CpG in the BRCA1 promoter versus WGS-derived tumor purity with pyrosequencing-derived methylation-state indicated in red. The FlexMix framework was initialized 10 times with the same parameters as those used for adjusting Illumina methylation beta values to identify subpopulations in the data, each iteration the identified populations were highly concordant with pyrosequencing-derived BRCA1 promoter methylation calls. **C)** Effect of increasing amount of noise on accuracy of FlexMix population calls versus pyrosequencing. Distributions derived from 50 repetitions for each noise level. **D)** As in (C) but showing the average beta-value shift introduced by purity-adjustment when increasing the amount of added noise.

## Discussion

Epigenetic alterations are a hallmark of cancer. Delineating such alterations may help in identification of genes important for tumorigenesis, but also aid in global characterization of epigenetic patterns associated with molecular subgroups of a malignancy. In most epigenetic analyses of bulk tumor tissue to date, the effect of tumor purity on methylation estimates has not been fully addressed despite that it has long been recognized that methylation estimates can be affected by differing epigenetic states present in the mixture of malignant and non-malignant cell types that make up the bulk tumor. Here, we present a simple strategy to adjust for this at an individual CpG level in high-dimensional DNA methylation data.

The concept of adjusting tumor methylation estimates using tumor purity estimates is not new and has been proposed for both bisulfite-sequencing [15] and Illumina array data [15,17,19–22]. Most notably among previous work directed at solving this issue for Illumina methylation data, the authors of the InfiniumPurify-package have developed several tools for addressing the issues raised in the current work. The most recent addition to the InfiniumPurify-package implements a mixture modeling approach to define differentially methylated (DM) CpGs between groups [22], while controlling for the influence of purity and between-group differences in methylation states. This approach however requires prespecified group-variables and does not produce “purified” methylation estimates for downstream use. The InfiniumPurify R-package does include a function for producing purified methylation estimates (function InfiniumPurify), however, this models the tumor compartment as one entity, leading to only minor improvements in analyses presented in our current investigation. In addition, the InfiniumPurify-package is currently only implemented for the 450K-platform. Other available methods have been developed in order to quantify cell type infiltration (e.g., Methylresolver [13] or epiDISH [10]) or perform differential methylation analysis between predefined groups and/or controlling for known confounding cell types (e.g., InfiniumDM [22] or CellDMC [11]). Taken together, we therefore believe there is still an unmet need to further develop flexible and generic algorithms that can address the issue of correcting tumor methylation estimates. To this end, we utilized automated population discovery using the FlexMix framework and linear regression-based adjustment of CpG methylation beta values. We show that our approach yields intuitively interpretable and biologically sound results when applied to a large cohort of breast cancer tumors collected by The Cancer Genome Atlas project. The main advantage of our method is an intuitive and simple approach, a deterministic output, and that the method is applicable cross-platform. We show that using a mixture modelling approach seems to outperform at least simple dichotomization as well as performing regression-based adjustment modelling of the tumor compartment as a single entity. Additionally, our approach can infer the “normal” background methylome with reasonable accuracy, eliminating the need for reference methylomes when a sufficiently large cohort is profiled. We believe that the general approach outlined in this work can be applied to most types of methylation data, potentially with purity variables obtained from many different sources, including the methylation array itself using tools included in packages such as InfiniumPurify [21] or MethylResolver [13]. This general framework can be improved in future work by e.g., refinement of the population discovery process and modelling of systematic bias affecting regression intercept terms and residuals. Moreover, considering the continuously growing body of public methylation profiles for different malignancies the concept of static reference sets usable for correcting small sized cohorts could be investigated. As currently devised, the method runs the full 450K array (630 samples x ~ 420 000 CpGs) in around 8 hours on a standard 4-core processor, and runtime scales linearly downward with a larger core/thread count. Notably, this work is not intended to produce an optimal method or perform extensive benchmarking against other methods, neither is it aimed at evaluating the feasibility of using different data sources for deriving input purity estimates. Instead, this work serves to illustrate a way of accomplishing the goal of reducing the impact of the non-tumor compartment on methylation estimates derived from the Illumina 450/850K arrays.

Our search for a method capable of adjusting methylation estimates to account for tumor purity started with our observations in the *BRCA1* gene locus [24]. Through this work it has become evident that the effect of non-tumor methylation is pervasive in bulk tumor data, and that thousands of loci are affected in any given high-throughput methylation experiment. The *RB1* gene represents another bona-fide tumor suppressor gene that has long been known to become inactivated through DNA methylation in cancer [25]. High impact genes such as *RB1* and *BRCA1* are epigenetically inactivated at high frequencies which is why they were detected in the early days of molecular cancer research. However, epigenetic silencing of key tumor suppressor genes may be infrequent but critical when appearing. An illustrative example is homologous recombination deficiency (HRD) in breast cancer. While promoter hypermethylation of *BRCA1* has been reported as the most frequent cause of HRD in TNBC, a substantial number of patients’ tumors still lack a known inactivation mechanism (driver alteration) [26]. In these tumors the HRD phenotype is likely conferred by alterations in less penetrant genes or through polygenic interactions, as illustrated by the infrequent (2%) promoter hypermethylation of *RAD51C* in TNBC that however confers a genetic HRD phenotype similar to *BRCA2* inactivation [26]. Improved processing of DNA methylation data can thus be a critical component in the search for novel but infrequently deactivated tumor suppressor genes. Another important issue made more addressable by our method is the question of which and how many pure epigenetic phenotypes exist in e.g., breast cancer, and what are the base epigenotypes on top of which the intrinsic subtypes reside. Epigenetic profiling of pure (or purified) methylomes may therefore provide novel insights into new drivers of HRD or improve our understanding of the base unconfounded tumor epigenome, allowing for improved patient stratification and understanding of the basic tumor biology of breast cancer.

## Conclusions

We present a conceptual method and an algorithm for correcting large-scale DNA methylation data for the influence of the TME on global methylation estimates. Our method uses a flexible and simple mixture modeling approach to identify tumor sample populations differentially influenced by non-tumor background and correct for this effect on a global basis. Our method also generates accurate estimates of the ground-state methylation of the non-tumor compartment and does not rely on available normal samples. We expect that our approach can be developed further and integrated into pre-existing methods to improve the capacity of these for de-noising bulk methylation data and allow for more unbiased epigenomic analyses. Additionally, we believe that in-depth analysis of loci that exhibit a *BRCA1*-like methylation pattern could yield novel leads in the search for tumor suppressor genes frequently inactivated by DNA methylation and that purified methylomes may provide valuable insights into the question of pure epigenotypes in cancer.

## Acknowledgments and Funding

The authors would like to thank Dr Dominik Glodzik at Harvard Medical School for suggesting the use of FlexMix as the basis for the proposed approach. We would also like to thank Dr Shamik Mitra at Lund University Department of Clinical Genetics for providing valuable input to the manuscript. We acknowledge the work of The Cancer Genome Atlas (TCGA) project and the National Cancer Institute Genomic Data Commons (NCI GDC) in making the underlying data available.

Financial support for this study was provided by the Swedish Cancer Society (JS: CAN 2021/1407 and a 2018 Senior Investigator Award, SIA190013), the Mrs Berta Kamprad Foundation (JS: FBKS-2020-5), the Swedish Research Council (JS: 2021-01800), and governmental funding (JS: ALF, grant 2018/40612). MA has received funding by The Gunnar Nilsson Foundation (GN-2018-5), and by the Mrs Berta Kamprad Foundation for cancer research (FBKS-21-27-328).

## Authors Contributions

*Conception and design*: Mattias Aine

*Collection and assembly of data*: Johan Staaf, Mattias Aine

*Data analysis and interpretation*: Johan Staaf, Mattias Aine

*Financial support:* Johan Staaf

*Manuscript writing:* All authors

*Final approval of manuscript:* All authors

*Agree to be accountable for all aspects of the work:* All authors

## Supporting Information

### Supporting Information 1

**Supplementary Figure 1:** Scaled density plot showing the correlation between individual sample beta values (N=630 samples) and average normal breast methylation (GSE67919) for unadjusted (black), purity adjusted (red), and inferred normal (orange) samples.

### Supporting Information 2

**Supplementary Figure 2:** Heatmap visualization of the top 5000 most varying CpGs in the TCGA BRCA data set in inferred normal (leftmost), unadjusted (left), InfiniumPurify-adjusted (right), and purity adjusted (rightmost) data.

### Supporting Information 3

**Supplementary Figure 3:** The beta distribution before and after purity-adjustment in distally located (>5kb from promoter) CpGs without overlapping ENCODE TF-sites stratified by the five hierarchical clustering subgroups (left), stratified by PAM50 subtypes (middle), stratified by TNBC-status (right)

